# BasecRAWller: Streaming Nanopore Basecalling Directly from Raw Signal

**DOI:** 10.1101/133058

**Authors:** Marcus Stoiber, James Brown

## Abstract

All current nanopore basecalling applications begin with the segmentation of raw signal into discrete events, which are ultimately processed into called bases. We propose the basecRAWller algorithm, a pair of unidirectional recurrent neural networks that enables the calling of DNA bases in real time directly from the rawest form of nanopore output. This shift in nanopore basecalling provides a number of advantages over current processing pipelines including: 1) streaming basecalling, 2) tunable ratio of insertions to deletions, and 3) potential for streaming detection of modified bases. Key to the streaming basecalling capability is sequence prediction at a delay of less than 1/100th of a second, allowing future signal to continuously modulate sequence prediction. BasecRAWller is computationally efficient enabling basecalling at speeds faster than current nanopore instrument measurement speeds on a single core. Further, basecalling can be paused and resumed without any change in the resulting predicted sequence, transforming the potential applications for dynamic read rejection capabilities. The basecRAWller algorithm provides an alternative approach to nanopore basecalling at comparable accuracy and provides the community with the capacity to train their own basecRAWller neural networks with minimal effort.

## Introduction

The Oxford Nanopore Technologies (ONT) MinION and other nanopore sequencing devices measure the nucleotide sequence of DNA (or RNA). This process begins by the measurement of electric current across a nanopore as strand of DNA passes through the pore. These measurements are stored as 16-bit integer data acquisition (DAC) values, currently taken at 4kHz frequency. With a DNA strand velocity of ∼450 base pairs per second, this gives approximately nine raw observations per base on average. This signal is then processed to identify breaks in the open pore signal corresponding to individual reads. These stretches of raw signal are basecalled --the process of converting DAC values into a sequence of DNA bases that is a prerequisite for essentially all applications of nanopore technology.

Extant basecalling algorithms, albacore (ONT), Nanocall(David, et al., 2017), and DeepNano(Vladimír Boža, 2016), all begin with the same initial processing step, that is to segment the DAC values into “events” that ideally represent individual nucleotides. These events are then processed with either a hidden markov model (HMM; Nanocall and previous versions of ONT basecallers) or a recurrent neural network (RNN; albacore and DeepNano) to predict the sequence of bases (k-mer) in and near the center of the pore associated with each event. These values are subsequently resolved with neighboring events to produce a complete sequence for each read.

We present the basecRAWller algorithm (available at basecRAWller.lbl.gov) to shift the nanopore basecalling paradigm by deferring segmentation into events until after sequence content is analyzed. By utilizing only unidirectional (forward in sequencing time) recurrent neural networks the basecRAWller algorithm predicts bases in a truly streaming fashion. Other extant basecallers require a full pass over the signal to produce final base calls. Single reads are approaching 1 MB(Miten Jain, 2017) and current DNA translocation speed of 450 bp/sec, so a single read can take over half an hour to sequence. Thus accurate streaming basecalling is incredibly beneficial for several applications, especially those related to dynamic read rejection(Loose, et al., 2016). We propose that fundamentally changing the way that nanopore reads are basecalled has substantial potential to increase the accuracy even further and open new possibilities for nanopore sequencing technology.

## Methods and Results

### basecRAWller Algorithm and Performance

The basecRAWller algorithm is composed of a pair of recurrent neural networks, referred to as the raw and fine-tune nets (Figure 1). The raw net is composed of several stacked long short term memory units (LSTMs; (Hochreiter and Schmidhuber, 1997)) followed by one or more fully connected layers. This network takes in scaled DAC values and outputs continuous multinomial logit predictions for each 4-mer associated with each observation. Importantly, these predictions are made at a delay of 33 raw observations (less than 1/100th of a second in sequencing time), which allows the network to take in signal at a brief lag past an observation before predicting the associated sequence allowing for modulation of the predicted sequence (Supp. Figure 1). This network simultaneously outputs the probability that each observation corresponds to the beginning of a new base and in the future could be modified to output additional factors including identification of modified bases. A segmentation algorithm is then applied to segment the continuous sequence predictions into discrete events. The 4-mer predictions, averaged over each event are then passed to the fine-tune net to produce final sequence predictions, including small inserted bases (2 bases per event) and event deletion. These predictions are made at a delay of two base pairs (approximately 0.005 seconds) and the output of this neural network comprises the final predicted sequence from the basecRAWller algorithm.

**Figure 1.**
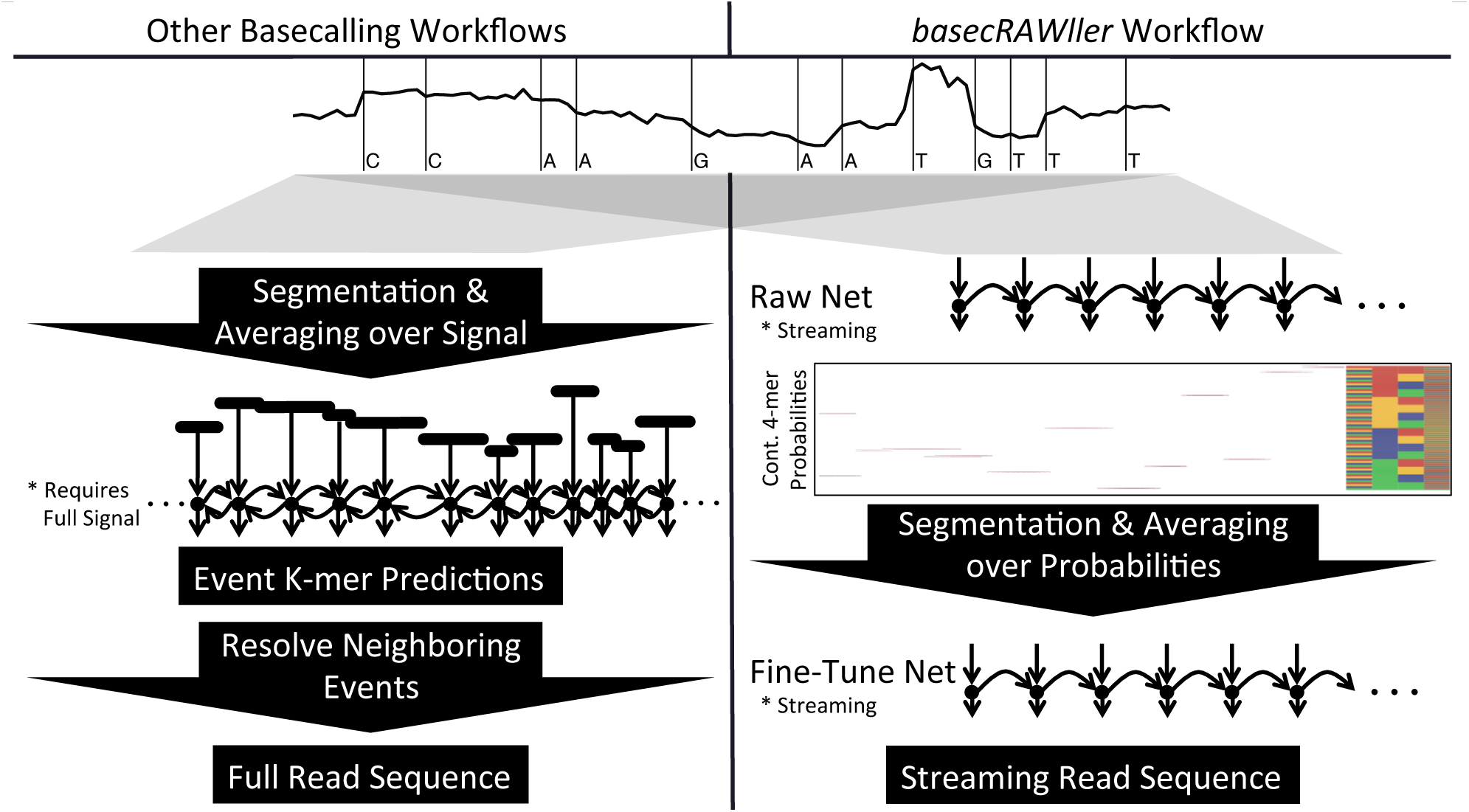
Comparison of current basecalling workflows with the basecRAWller workflow. Key differences allow for streaming basecalling using the basecRAWller algorithm that is not achievable with current algorithms based on the initial segmentation step.

We have applied the basecRAWller algorithm to two data sets from: E. coli (two control samples from (Marcus H Stoiber, 2016)) and Human (Nanopore WGS Consortium(Miten Jain, 2017); NA12878). These samples were processed with entirely different procedures: different library preparation workflow (FLOMIN-105 vs 1D ligation), translocation speed (250 bps vs 450 bps), read types (1D vs 2D) and flow cell (R9 vs R9.4) so these data sets represent quite distinct raw nanopore data. Performance of neural networks trained and tested on either data set (Figure 2) reveal that a basecRAWller model trained on the human data set is more robust across the two diverse data production pipelines, most likely due to the DNA translocation speed which significantly enhances challenges associated with sparse data at individual bases. We note that the basecRAWller model trained on the E. coli data set is robust to an independent held out E. coli replicate (different library preparation and flowcell; same lab). Moving forward, models may be trained based on each version of the nanopore technology or sample type and applied where applicable (including translocation speed, native versus amplified, ratchet protein and other workflow difference). Models can easily be shared amongst the community enabled by the basecRAWller framework. Additionally, particular parameters can be tuned to alter error profiles. For example the insertion penalty training parameter produces basecRAWller models with a range of insertion to deletion ratios (Supp. Figure 2). A full description of basecRAWller training parameters can be found in the supplementary methods and Supp. Table 1. Currently (basecRAWller version 0.1) the model trained with the human data is the default provided with for the basecRAWller software.

**Figure 2.**
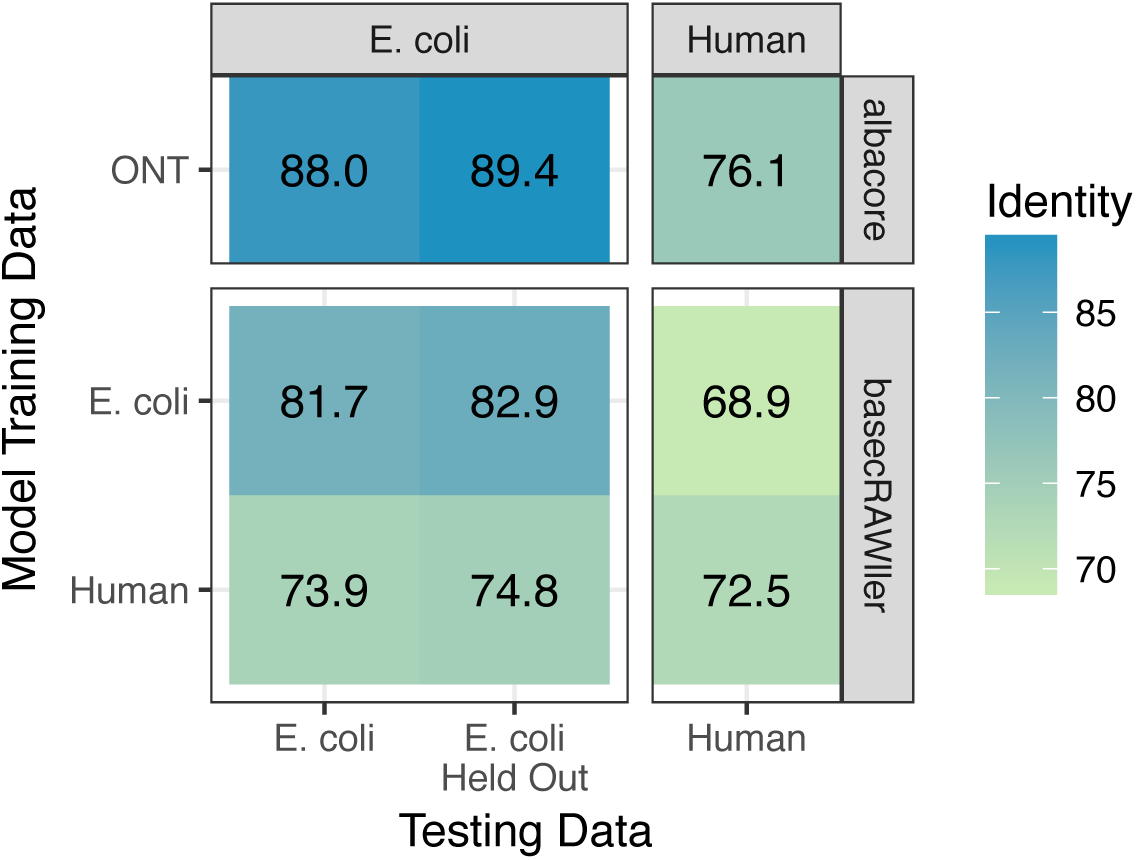
BasecRAWller performance. Data used for testing on the x-axis and data used for training on the y-axis. Oxford Nanopore Technologies (ONT) basecaller albacore (based on unknown data) in the top panel is compared to the basecRAWller trained models.

Comparison to basecalls from Oxford Nanopore Technology’s (ONT) albacore basecaller shows that basecRAWller is performing worse (Figure 2), paying a penalty for the advantage of streaming performance. Further, we note that there is room for improvement with additional training and basecRAWller neural network hyper-parameter selection --both tasks are enabled by the basecRAWller software package. We note that all hyper-parameter testing was completed using the E. coli data set, so it is likely that increased performance will be achieved in the human data set with additional hyper-parameter searching.

### basecRAWller Runtime Performance

Essential for streaming performance is the computational efficiency of basecRAWller – allowing basecalling at speeds faster than current nanopore instrument measurement speeds. BasecRAWller neural network models are based on the tensorflow python software interface(Martín Abadi, et al., 2015). On a single Intel Xeon E5-2698 v3 (2.3GHz) processor we achieve read processing speed of greater than 8000 observations per second, twice as fast as current sequencer data collection speeds. Processing is concentrated on the raw net, which consumes ∼90% of the processing time, while the segmentation algorithm takes a negligible amount of time and the fine-tune net consumes the remaining 10% of processing time. In future iterations of basecRAWller models it is of note that some parameters are sensitive to processing time. In particular the number and size of LSTM layers used in the raw net have a large effect on the final processing speed of a basecRAWller model.

### DAC Signal Normalization

The basecRAWller algorithm begins from a contiguous section of nanopore data acquisition (DAC) values that corresponds to a continuous strand of DNA. As there is drift in the relative levels of DAC values the first step in the basecRAWller algorithm is to normalize the DAC values. Median normalization, as reported in Stoiber *et al*.(Marcus H Stoiber, 2016) and employed in the current basecRAWller version requires the full read signal (or some proportion thereof) and is thus inhibitory to streaming capabilities. To overcome this roadblock, we investigated the signal immediately upstream of the read start and found that this signal level correlates very well with the median signal within the read (r ∼ 0.76) and could thus be used as a surrogate for streaming applications (Figure 3A). We note that we are currently using a simplistic algorithm to identify signal stretches corresponding to initially occupied pores, and hence this correlation could be made substantially stronger with more accurate identification procedures (Figure 3B-C). Leveraging before read start median normalization, the basecRAWller pipeline comprises the first fully streaming nanopore basecalling application.

**Figure 3.**
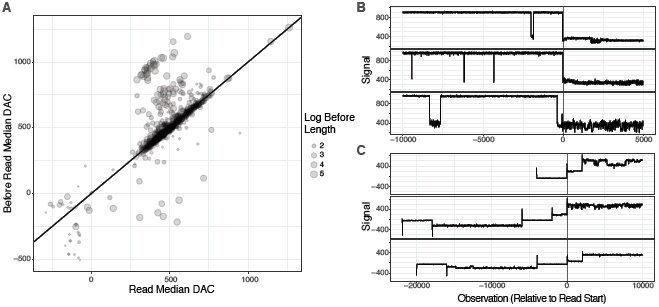
Enabling full streaming pipeline by using median of nanopore signal from before the beginning of a read. **A.** The correlation between the median signal level before a read and within a nanopore read from human data. **B.** Examples of several reads with before read signal levels significantly higher than within read signal levels. For these reads the end of the open pore is identified incorrectly based on small pore blockages and thus the before read signal includes stretches of open pore signal. **C.** Examples of several reads with before read signal levels significantly lower than within read levels. These are likely spuriously identified reads associated with pore clearing voltage flicks and should be removed in future algorithm versions.

## Discussion

The basecRAWller algorithm, along with the previously published nanoraw software package, provides an alternative approach to nanopore basecalling, which potentiates real-time nanopore sequencing and enables the community to train basecRAWller neural networks with a felicitous command line API. The neural networks fitted to date required ∼250,000 compute hours on Intel Xeon processors at LBNL NERSC computing facility. We anticipate continued improvement of models with improved hyper-parameter optimization, as well as inclusion of community involvement in basecRAWller improvements.

An important future direction for the basecRAWller pipeline will be adding the capacity to detect, in real time, covalently modified bases. Such capabilities have the potential to be transformative for cancer and toxicological research, and more broadly for foundational and environmental biology. Just as base start positions are an output of the raw net, modified bases can be identified without combinatorial expansion in the space of 4-mers. Additionally, basecalling directly from raw signal could be performed with a bi-directional neural network and may show significant increases in prediction accuracy over extant basecallers, though obviously removing the streaming capacity of basecRAWller. The basecRAWller software minimizes barriers to entry for the community to begin, much more broadly, to develop diverse nanopore basecalling algorithms, and, ultimately, to explore the space of extant DNA modifications throughout the biosciences.

## Author Information

M.H.S. has received travel expenses and accommodation from Oxford Nanopore Technologies to speak at organized symposia.

## Supplemental Methods

### Raw Neural Network

Once normalized DAC values have been computed, they are fed into a recurrent neural network (RNN), specifically a number of unidirectional (in the direction of sequencing time) long short term memory (LSTM) layers followed by one or more fully connected layer. The structure of this first neural network (termed the raw net) is depicted in Supp. Figure 3. This network takes as input a single scaled DAC value and returns the probability that each 4-mer is represented by the appropriately offset DAC value (represented by logit values), as well a “memory” that is passed on to the next observation along with the next scaled DAC value. In parallel to sequence output, the raw net produces the probability that the appropriately offset position is the beginning of a new base to be potentially used in the segmentation step. This network can be constructed with any number of LSTM layers each with any number of output units (entered as a parameter during training). The 4-mer to predict is also variable by specifying the number of bases before and after the current base to be included. For all analyses presented here we use a three-layer network with 75, 100 and 50 output nodes and output 4-mers. The third layer is added after training the two-layer network in order to more quickly identify appropriate parameters for the first two LSTM layers. The basecRAWller software provides this functionality.

Constructing appropriate training data sets and setting training parameters is key to the success of basecalling algorithms. Starting with basecalled reads by the using the ONT software, we construct training data by applying the *nanoraw* software package(Marcus H Stoiber, 2016) genome_resquiggle algorithm. This algorithm resolves error prone base calls with a genomic alignment so that raw signal is assigned to potentially perfect basecalls from a known source (here E. coli or human chromosome 21). For the raw net this allows the assignment of each raw DAC value with the true 4-mer centered on that position and defined by a number of positions before and after the current position.

To construct training and testing data sets, nanopore reads are provided (7,500 are provided for both presented trained models). For each read, a random starting position between the start and middle of a read is selected. This is to avoid any over-fitting to the start of reads that would all be presented at once to be trained in the initial iterations of each epoch. Only reads longer than the 50^th^ percentile of read lengths are included in the training data (3,740 reads included for fitted models; this value showed good robustness with held out test data with reasonable computational efficiency). This is so that a training epoch (all provided data) is long enough to allow inclusion of sufficient variety and quantity of data to remediate over-fitting early in the training process. These reads each starting at their randomly selected position are combined into a matrix with the length of the 50^th^ percentile read length (i.e. the shortest read included in the training data set). Additionally, another vector is provided with the true start positions for each base in each read to train the base start raw net output. This procedure is encoded in the prep_raw_net command in the basecRAWller software package.

To more accurately predict sequence from raw signal, the raw net is trained to predict the sequence at a delay. For both networks presented here this delay is set to 33 observations (less than 1/100th of a second in sequencing time), which was chosen after testing many values in this range (Supp. Figure 1). This allows the network to take in the signal well past a given observation, which modulates the predicted sequence, before producing a final sequence prediction. This parameter shows similar performance (data not shows) within the human training data, despite the different translocation speeds.

In addition to these high-impact parameters there are several parameters specific to the raw net training (all parameters are included in Supp. Table 1). The following parameters were tested for effect on model accuracy as well as computational training time. The number of observations included in each training iteration (number of unrolled observations) has a reasonable effect on the final output with values greater than 300 showing reasonable results. The value used for both trained models was 500 as greater values were again prohibitively computationally expensive. The neural network learning rate was set to 1.0 for presented trained models, with values between 0.5 and 1.5 showing reasonable results. In order to stabilize the model, as it gets closer to a global minimum loss, the learning rate is halved every 500 observations. This value showed reasonable results in the range of 200-10000 observations. The momentum parameter is used by the gradient ascent algorithm to maintain the direction of fitting from iteration to iteration and was set to 0.9 for both models presented here. Values between 0.85 and 0.99 show reasonable results. Given the outputs of sequence probabilities and base start positions, a parameter is provided to control the proportion of the loss attributed to the base starts cross-entropy loss and the proportion attributed to the k-mer sequence cross-entropy loss. This parameter is set to 1.1 (10% more weight attributed to start position loss) for both fitted models. Values between 0.8 and 1.5 show reasonable results. In order to allow the base start probability predictions to “ramp up” before and “ramp down” after a true base start, positions within a range around true base starts are masked (their loss values are set to 0). For both fitted models two observations around the true starts were masked, with reasonable values being between 0 and 3. During training the probability that any connection between two nodes is included is available as a parameter to basecRAWller, but this parameter showed little improvement in the fitted model, so all connections are included during training (parameter value 1.0).

The raw neural network is also designed to output events back into the raw files thus allowing iterative training of basecRAWller models (i.e. preparing training data for a basecRAWller model from a basecRAWller trained basecaller). We have found limited success with this procedure thus far, but make this capacity available to the community and plan to include this as an output from the fine-tune net soon.

In addition, the capacity to restore the training of the raw net with a new LSTM layer (or with a new fully connected layer or hyper-parameters) is provided by the basecRAWller software. This restoration process restores the trained parameter values to the LSTM layers, adds a new LSTM layer with a specified node size and fully connected layers that are not restored due to the new fully connected layer that connects to the new LSTM layer. This procedure was employed during the optimization of the number of LSTM layers for the raw trained nets in this manuscript.

### Segmentation

The raw net produces, for each raw observation, the probability that the appropriately offset observation is derived from each distinct 4-mer, which then needs to be segmented into predicted bases. The raw net also produces the probability that each position is the start of a new base. For the basecRAWller models here we have used these raw net base start predictions for segmentation, but we propose two other measures (running Hellinger distance and entropy difference) that may perform better in particular settings.

For any of these measures, peaks are identified. Peaks are required to be 4 observations apart from another peak, but this value is a tunable parameter. To save computational time during training, only the first 100,000 observations (25 seconds) are used for segmentation metric computations. From these valid peak locations we define a global cutoff value and take all peaks with value above this threshold as the defining points for segmentation into events.

A command in the basecRAWller software “analyze_splitting” is provided in order to identify an optimal cutoff for any trained networks. This optimal value is determined by analyzing some number of reads and choosing the global cutoff value that minimizes the difference between the true number of bases for each read and the number of identified segments given a global cutoff value. This function can also test all three measures and select the measure and cutoff value based on relative performance (default). This value and the selected measure is stored within the graph file for fine-tune net training and final basecalling applications.

Alternative measures, Hellinger distance and entropy difference, are computed from the matrix of 4-mer logits at some fixed offset before and after each observation. Hellinger distance is a natural measure of the distance between two multinomial distributions, which is the exact realm we find when looking for segmentation locations. It is natural for the probability distribution to shift considerably centered at a transition between true bases. Similarly, entropy is a measure of a statistical distribution and shifts in entropy through time again indicate a shift from one base to another. Upon selection of an alternative measure, the fine-tune net needs to be re-trained.

After segmentation positions are identified, a matrix is prepared in order to train the fine-tune net or dynamically for basecalling. For each read the raw net 4-mer probability distribution is averaged within each segment to be passed to the fine-tune net.

### Fine Tuning Neural Network

After raw net processing, segmentation and distribution averaging a second neural network is applied to fine-tune the predicted bases from the raw net. The structure of the fine-tune net is identical to the raw net with some number of LSTM layers followed by one or more fully connected layer that outputs sequence probability values. The difference here is that each observation taken in by the fine-tune net is a vector of logits for each possible k-mer, instead of a single normalized DAC value and the output is zero, one, or more bases represented by this segment. For analysis presented here we use two LSTM layers with 200 and 75 nodes. The output of the fine-tune net is the logit values for between 0 and some length of k-mer that represent the bases to assign to the current segment. The 0-length state indicates that the current segment should be deleted and should have been joined with the previous segment. The 1-mer states indicate that this segment was correctly assigned a single base and specifies the specific base for that segment. 2-mer and greater states indicate that the current segment represents more than one base (was under segmented). The maximal number of inserted bases for all analyses presented here is set to two, but this is again a parameter for tuning future basecRAWller models available in the basecRAWller software.

As with the raw net, the fine-tune net is trained to predict bases at a delay in order to take in values past the current base allowing for modulation of the sequence predictions given values slightly ahead of the current position. For both models presented here this offset was set to 2 segments, representing approximately 18 observations or again less than 1/100th of a second. This parameter has a smaller, but discernible effect than the raw offset parameter.

The construction of the training data for the fine-tune net is key as there is now a discordance between the identified segmentation positions from the raw net (raw segments) and the ideally true segments identified by the nanoraw genome_resquiggle assignments (true segments). Raw segment are assigned a true base as long as the true segmentation position is not more than 3 positions past the raw segment position. Any true base included within the raw segment but more than 3 positions into the current raw segment will be included in the next raw segment (inserted bases). If more than 2 extra true bases are assigned to a raw segment, the assigned true bases are trimmed to include only the 3 most recent bases. If no true segment exists between the last raw segment position and the current one, then the raw segment is assigned no bases (deleted event). Given this training setup the output of the fine-tune net will represent the final sequence for a read provided in a completely streaming fashion.

Given this output scheme for the fine-tune net, the deletion categories are prone to over-prediction. Any deleted base is represented by a single deletion category, while a single inserted base is represented by 16 output categories (for all combinations of the assigned base and the inserted base). Thus a parameter is provided to maintain equal (or tune-able) insertion to deletion rates. In order to achieve approximately equal insertion and deletion rate this parameter is set to 2.0. During training, deleted event loss values are deflated by a factor of 2-1=0.5, single base insertion categories loss values are inflated by a factor of 2^1^=2 and two base insertion category loss values are inflated by a factor of 2^2^=4. Tuning this parameter allows basecRAWller models with different insertion to deletion ratios, which may be of value in particular situations.

As with the raw net there are several general neural network training parameters which need to be set. The number of observations included in fine-tune net training is set to 90 for models presented here, with values around 100 showing reasonable results. The learning rate halving parameter is set to 3,000 for the fine-tune net. Additional, parameters are set very similarly to the raw net: learning rate is set to 0.9 and momentum is set to 0.9.

## Supplemental Table

**Supplementary Table 1.** Description of all parameters used in training raw and fine-tune nets along with intuition for impact on resulting performance, selected value, recommended ranges and general notes.

| Parameter | Training Step | Impact | Selected Value | Recommended Range | Related Parameters | Notes |
| --- | --- | --- | --- | --- | --- | --- |
| **Before Bases** | Raw Net | + | 1 | 1-4 | After Bases | Increasing the k-mer size will add many categorical outputs and may have deleterious effects above 4-mers |
| **After Bases** | Raw Net | + | 2 | 1-6 | Before Bases |  |
| **Observations Offset** | Raw Net | ++++ | 33 | 10-50 | Unrolled Observations | Probably the most important parameter in the whole pipeline |
| **Number of Reads** | Raw Net | ++ | 10000 | >5000 | Trim Percent | Important for a robust model; compare train and test accuracy for low values |
| **Trim Percent** | Raw Net | - | 50 | >10 | Number of Reads | This parameter ensures epoch lengths that are long enough; enough distinct training batches to avoid over fitting |
| **LSTM Sizes** | Raw Net | +++ | 75, 100, 50 | >25 | Fully Connected Sizes | More layers and larger sizes are better, but hurt final computational efficiency; Large and many layers may have trouble fitting well |
| **Fully Connected Sizes** | Raw Net | + | NA | >25 | LSTM Sizes, K-mer Bases | Not thoroughly tested; may increase computational speed, while maintaining accuracy |
| **Unrolled Observations** | Raw Net | +++ | 500 | >150 | K-mer Bases, Fully Connected Sizes, LSTM Sizes | LSTM layers can only "communicate" effectively over this range; May be strongly effected by chemistry version |
| **Learning Rate** | Raw Net | ++ | 1 | 0.5-1.5 |  | Very high values compared to many other NN learning problems |
| **Learning Rate Half Life** | Raw Net | + | 500 | 200-2000 |  | Adjust to fine tune the network once global minimums are approached |
| **Momentum** | Raw Net | - | 0.9 | 0.8-0.96 |  |  |
| **Total Iterations** | Raw Net | + | 5000 | >2000 |  | Very important parameter for model robustness |
| **Start Proportion** | Raw Net | ++ | 1.1 | 0.7-1.5 | K-mer Bases, Learning Rate | Low start accuracy produced by lower values may not produce worse results in splitting algorithm, so don't completely depend on the start accuracy for performance of this parameter |
| **Start Mask Size** | Raw Net | + | 2 | 0-3 |  | Allows for start probability to "ramp up" and "ramp down" around the true start position |
| **Keep Probability** | Raw Net | - | 1 | 0.9-1.0 |  | Does not seem to help robustness; set to 1 |
| **Measure Lag** | Splitting | - | NA | 1-3 |  | Only applies to Hellinger and Entropy Distances; LSTM usually performs best |
| **Minimum Observations per Base** | Splitting | - | 4 | 2-6 | Measure Lag |  |
| **Number of Split Reads** | Splitting | - | 5000 | >1500 |  | More is better; more compute time for more reads though |
| **Max Observations per read** | Splitting | - | 100000 | >20000 |  | Helps keep compute time reasonable for very long reads |
| **Maximum Inserted Bases** | Fine-Tune Net | + | 2 | 1-4 | K-mer Bases, Events Offset | As with k-mer bases large sizes will cause many categorical outputs and may deleteriously effect performance |
| **True Split Offset** | Fine-Tune Net | + | 3 | 1-6 | Maximum Inserted Bases | This is the wiggle room to let a true offset be assigned to a discovered segment when they are very close; it should probably not be set larger than Minimum Observations per base, but this has not been tested |
| **Events Offset** | Fine-Tune Net | ++ | 2 | 0-5 | Learning Rate | Similar to Observation offset parameter, but seems to have a smaller effect |
| **Insertion Penalty** | Fine-Tune Net | +++ | 2 | 1.0-3.5 |  | Training statistics are not comparable across values; Assess by final mapping percent identity |
| **LSTM Sizes** | Fine-Tune Net | +++ | 200, 75 | >50 | Fully Connected Sizes, K-mer Bases | More layers and larger sizes are better, but hurt final computational efficiency; Large and many layers may have trouble fitting well |
| **Fully Connected Sizes** | Fine-Tune Net | + | NA | >25 | LSTM Sizes, Maximum Inserted Bases | Not thoroughly tested; may increase computational speed, while maintaining accuracy |
| **Number of Unrolled Events** | Fine-Tune Net | ++ | 90 | >75 | Events Offset | LSTM layers can only "communicate" effectively over this range; May be strongly effected by chemistry version |
| **Learning Rate** | Fine-Tune Net | ++ | 0.9 | 0.5-1.2 |  | Very high values compared to many other NN learning problems |
| **Learning Rate Half Life** | Fine-Tune Net | + | 3000 | >1000 |  | Adjust to fine tune the network once global minimums are approached |
| **Momentum** | Fine-Tune Net | - | 0.9 | 0.8-0.96 |  |  |
| **Total Iterations** | Fine-Tune Net | + | 10000 | >5000 |  | Very important parameter for model robustness |
| **Keep Probability** | Fine-Tune Net | - | 1 | 0.9-1.0 |  | Does not seem to help robustness; set to 1 |

## Supplemental Figures

**Supplemental Figure 1.**
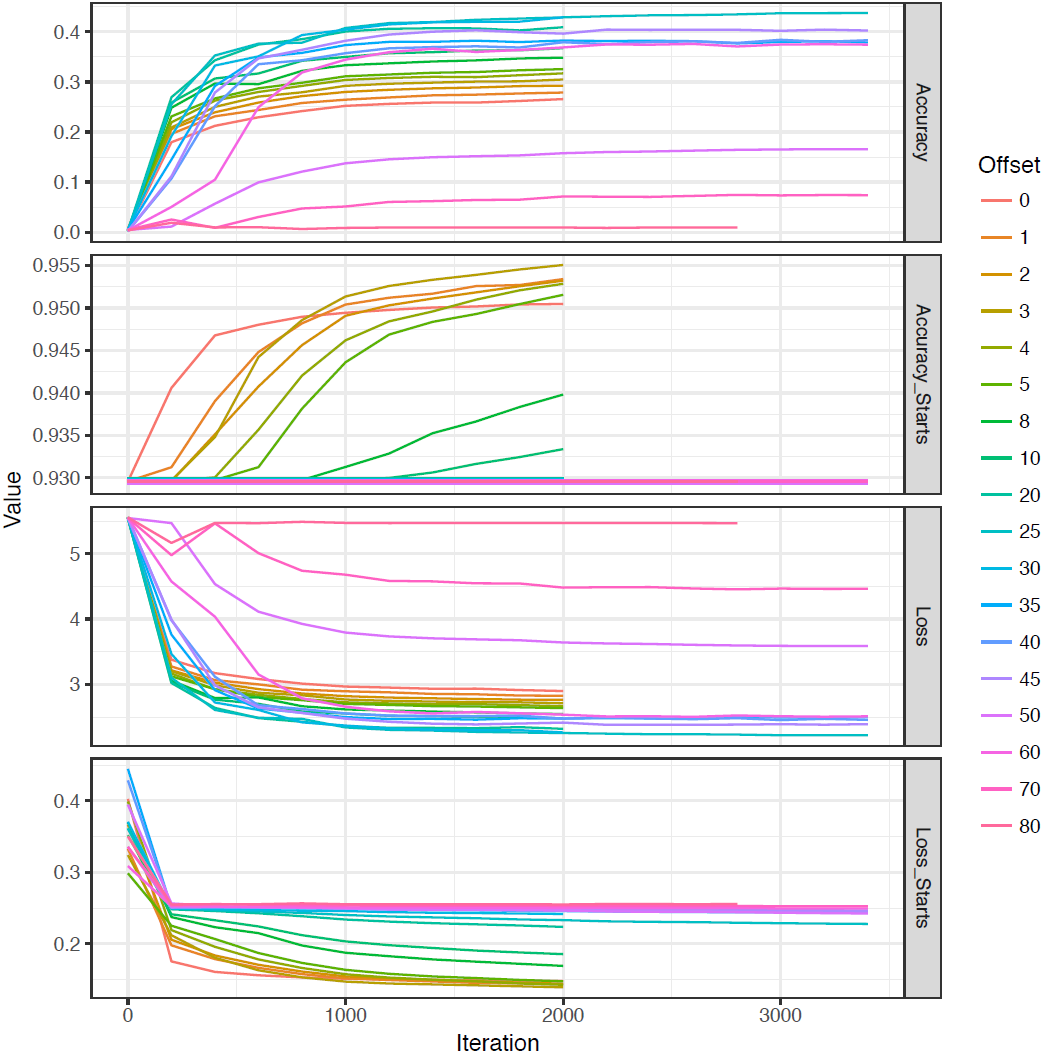
Raw net fitting performance over iterations for a range of the offset parameters. The final chosen value for trained nets was 33. Additional parameters were tuned to improve start location predictions, which show poor performance here.

**Supplemental Figure 2.**
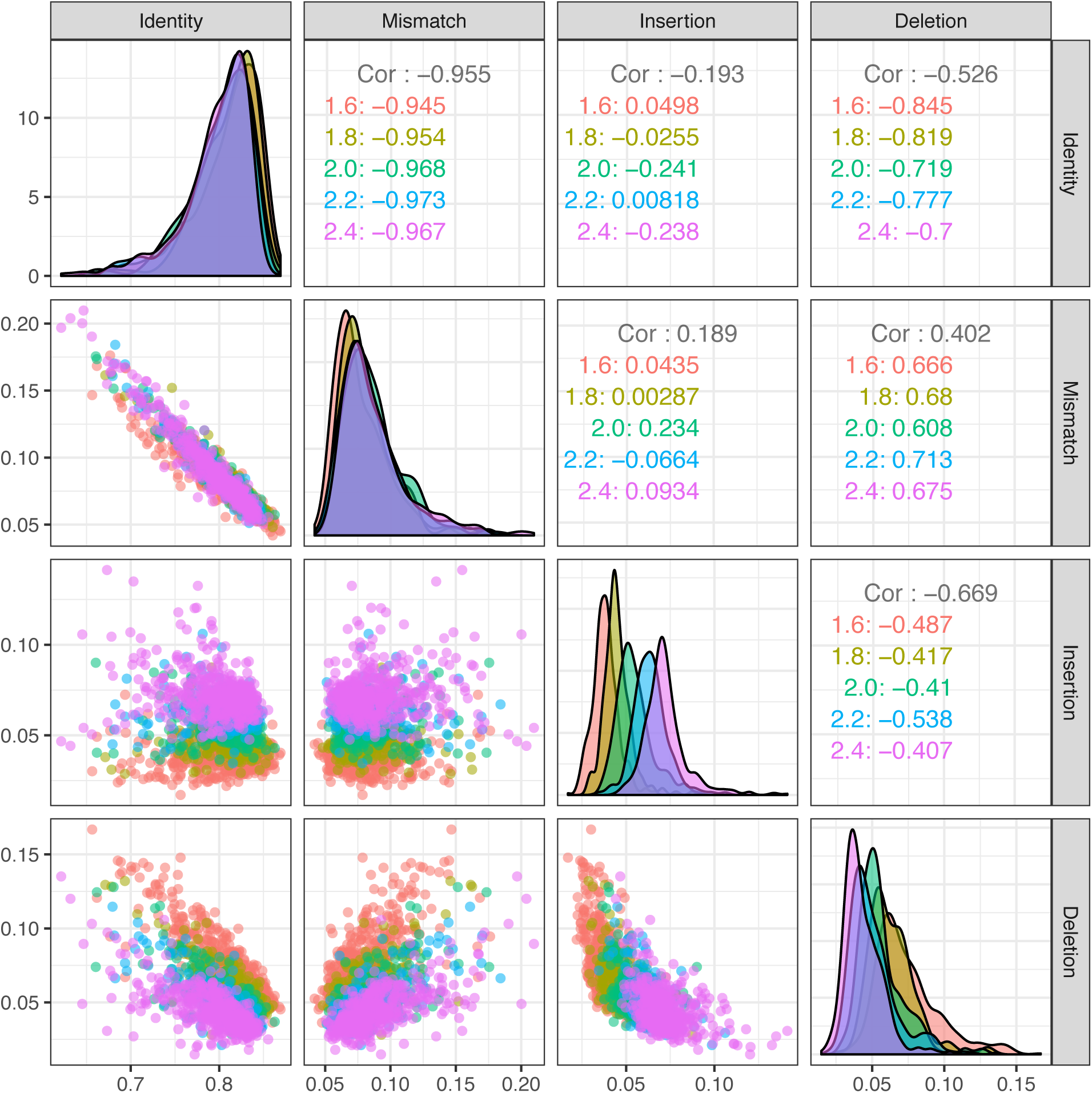
Comparison of mapping statistics for over 5 models trained with the same parameters except for the insertion penalty training parameter. Models show similar percent identity and percent mismatch while showing a range of insertion and deletion rates.

**Supplemental Figure 3.**
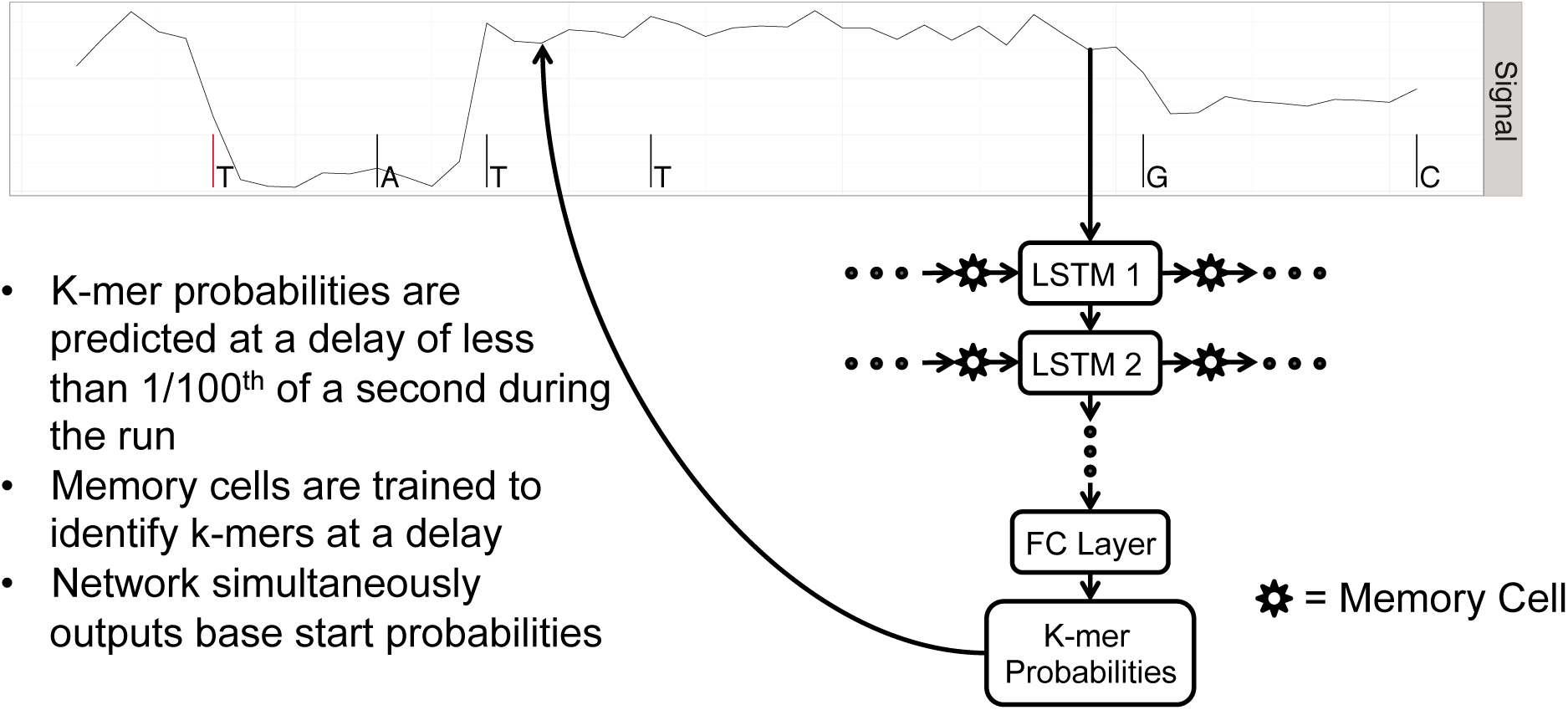
Raw and fine-tune neural network structures. The raw net is composed of three LSTM layers and one fully connected (FC) layers and the fine-tune net is composed of two LSTM layers and one FC layer. As noted probabilities are made at a delay in both nets and the raw net simultaneously predicts the probability of a base starting at that observation. The input to the fine-tune net is the segmented, mean output logits from the raw net and the output of the fine-tune net are the final sequences represented by each event.

## References

David, M., et al. Nanocall: an open source basecaller for Oxford Nanopore sequencing data. Bioinformatics 2017;33(1):49–55.

Hochreiter, S. and Schmidhuber, J. Long short-term memory. Neural Comput 1997;9(8):1735–1780.

Loose, M., Malla, S. and Stout, M. Real-time selective sequencing using nanopore technology. Nat Methods 2016;13(9):751–754.

Marcus H Stoiber, J.Q., Rob Egan, Ji Eun Lee, Susan E Celniker, Robert Neely, Nicholas Loman, Len Pennacchio, James B Brown. De novo Identification of DNA Modifications Enabled by Genome-Guided Nanopore Signal Processing. bioArxiv 2016;094672.

Martín Abadi, A.A., Paul Barham, Eugene Brevdo,, et al. TensorFlow: Large-scale machine learning on heterogeneous systems. 2015.

Miten Jain, S.K., Josh Quick, Arthur C Rand, Thomas A Sasani, John R Tyson, Andrew D Beggs, Alexander T Dilthey, Ian T Fiddes, Sunir Malla, Hannah Marriott, Karen H Miga, Tom Nieto, Justin O’Grady, Hugh E Olsen, Brent S Pedersen, Arang Rhie, Hollian Richardson, Aaron Quinlan, Terrance P Snutch, Louise Tee, Benedict Paten, Adam M. Phillippy, Jared T Simpson, Nicholas James Loman, Matthew Loose. Nanopore sequencing and assembly of a human genome with ultra-long reads. bioaRxiv 2017.

Vladimír Boža, B.B., Tomáš Vinar. DeepNano: Deep Recurrent Neural Networks for Base Calling in MinION Nanopore Reads. arXiv 2016.

